# Connectional architecture of a mouse hypothalamic circuit node controlling social behavior

**DOI:** 10.1101/445312

**Authors:** Liching Lo, Dong-Wook Kim, Shenqin Yao, Ali Cetin, Julie Harris, Hongkui Zeng, David J. Anderson, Brandon Weissbourd

## Abstract

Type 1 Estrogen receptor-expressing neurons in the ventrolateral subdivision of the ventromedial hypothalamus (VMHvl^Esr1^) play a causal role in the control of social behaviors including aggression. Here we use six different viral-genetic tracing methods to map the connectional architecture of VMHvl^Esr1^ neurons. These data reveal a high level of input convergence and output divergence (“fan-in/fan-out”) from and to over 30 distinct brain regions, with a high degree (~90%) of recurrence. Unlike GABAergic populations in other hypothalamic nuclei controlling feeding and parenting behavior, VMHvl^Esr1^ glutamatergic neurons collateralize to multiple targets. However, we identify two anatomically distinct subpopulations with anterior vs. posterior biases in their collateralization patterns. Surprisingly, these two subpopulations receive indistinguishable inputs. These studies suggest an overall system architecture in which an anatomically feed-forward sensory-to-motor processing stream is integrated with a dense, highly recurrent central processing circuit. This architecture differs from the “brain-inspired” feed-forward circuits used in certain types of artificial intelligence networks.

**SIGNIFICANCE:** How the cellular heterogeneity of brain nuclei maps onto circuit connectivity, the relationship of this anatomical mapping to behavioral function, and whether there are general principles underlying this relationship, remains poorly understood. Here we systematically map the connectivity of estrogen receptor-1-expressing neurons in the ventromedial hypothalamus (VMHvl^Esr1^), which control aggression and other social behaviors. We find that a relatively sparse, anatomically feed-forward sensory-to-motor processing stream is integrated with a dense, highly recurrent central processing circuit. Further, the VMHvl contains at least two subpopulations of Esr1^+^ neurons with different cell body characteristics and locations, with distinct patterns of collateralization to downstream targets. Nevertheless, these projection-defined subpopulations receive similar inputs. This input-output organization appears distinct from those described in other hypothalamic nuclei.

## INTRODUCTION

The hypothalamus controls an array of innate survival behaviors and associated internal motivational states, including feeding, drinking, predator defense and social behaviors such as mating and fighting (1, 2). It is comprised of multiple nuclei, each of which is thought to play a role in one or more behavioral functions. The connectivity of these nuclei has been extensively mapped using classical anterograde and retrograde tracing techniques (reviewed in (3, 4), revealing complex inputs to and outputs from these structures, as well as dense interconnectivity between them (5). It is increasingly clear, however, that hypothalamic nuclei are both functionally (6, 7) and cellularly (8, 9) heterogeneous. How this cellular heterogeneity maps onto connectivity, the relationship of this anatomical mapping to behavioral function, and whether there are any general principles underlying this relationship, remains poorly understood.

Genetic targeting of neuronal subpopulations and viral tools for tracing neuronal connectivity (10, 11), have provided a powerful approach to this problem in other brain systems (12-19). In the hypothalamus, viral tracers have been used to map the outputs of Agrp+ neurons in the arcuate (ARC) nucleus, which control the motivation to eat (20, 21). Although the ARC as a whole projects to ~6 different regions, these studies suggested that different subsets of ARC^Agrp^ neurons project to different targets (22). Similarly, studies of Galanin-expressing neurons in the medial preoptic area (MPOA) (23) have suggested that different MPOA^GAl^ neurons controlling different functions (pup-gathering, reward, etc.) project to distinct downstream targets (24) with little to no collateralization of projections.

Together, these and other data (25) have led to a view in which the hypothalamus consists of multiple “labeled lines” of genetically and functionally distinct neuronal subpopulations that map to anatomically distinct pathways. This stands in contrast to viral-anatomical studies of monoamine neuromodulatory systems: in the Locus Coereleus, neurons collateralize extensively and receive similar input regardless of downstream target (13), while in the Dorsal Raphe, subpopulations collateralize broadly but to distinct combinations of targets, and receive biased inputs depending on their projection target (26).

Type 1 estrogen receptor (Esr1)-expressing neurons in the ventromedial hypothalamic nucleus (VMH) control social and defensive behaviors as well as metabolic states (6, 7, 9, 27); reviewed in (28-33). Unlike the ARC^Agrp^ and the MPOA^Gal^ populations, which are primarily GABAergic (24, 34), VMHvl^Esr1^ neurons are mainly glutamatergic (35). Inputs and outputs of the VMHvl and the anatomically overlapping hypothalamic attack area (HAA) (36) have been extensively mapped in rats, using conventional anterograde and retrograde tracers (37, 38). In mice, one study using a genomically integrated axonal tracer, PLAP, mapped a subset of projections from progesterone receptor-expressing VMHvl neurons, which are highly overlapping with VMHvl^Esr1^ neurons (27). However, inputs to and projections from VMHvl^Esr1^ neurons, the extent of their collateralization, and the relationships between specific inputs and outputs have not been systematically investigated.

In this study we have used six different genetically targeted viral tracing methods to map the meso-scale connectivity of VMHvl^Esr1^ neurons. Collectively, these methods reveal that these neurons display a high degree of convergence, divergence, and recurrence in their inputs and outputs, respectively. However, unlike ARC^Agrp^ and MPOA^Gal^ GABAergic neurons, VMHvl^Esr1^ neurons collateralize to multiple downstream targets. Nevertheless, we identify two novel, non-overlapping Esr1^+^ neuronal subsets with anatomically distinct patterns of collateralization. Together, these data suggest a system architecture in which VMHvl^Esr1^ neurons receive sparse input from anteriorly located sensory structures and relay this information through two divergent processing streams: a sparse feed-forward relay targeting posterior (premotor) regions; and a highly recurrent amygdalo-hypothalamic network that may control decision-making and/or internal states.

## RESULTS

### Extensive recurrent connectivity of projections and inputs of VMHvl^Esr1^ neurons

Inputs to and projections from the VMHvl have been mapped in several studies using classical techniques in rodents (e.g. CTb and PHAL, (36-44). However, these techniques are unable to distinguish genetically-defined cell populations within VMHvl, which is heterogeneous (9). Further, with a few exceptions (45, 46) most studies report on a single type of tracing method (e.g., anterograde or retrograde), making it difficult to directly compare results across publications. To compare more directly the inputs and outputs of VMHvl^Esr1^ neurons, we used *Esr1-Cre* knock-in mice (7), together with Cre-dependent anterograde (12) and modified monosynaptic retrograde rabies (10, 11) viruses (see Methods), stereotaxically injected into VMHvl, and analyzed the results using serial 2-photon tomography (Tissue-Cyte, (47) at 100 μm z-intervals with 0.35 μm x-y resolution. Esr1^+^ neurons form a continuous population that extends from the VMHvl into the neighboring Tuberal nucleus (TU). Across experiments described in this study, the majority (~77%) of starter cells were located in these two regions combined, a population referred to hereafter as VMHvl^Esr1^.

Anterograde (non-transneuronal) tracing using Cre-dependent AAV-GFP (12) revealed projections of VMHvl^Esr1^ neurons to over 60 target regions (Fig. 1A-F, Fig. S1A, see Table 1 for brain region abbreviations). Most of these targets were previously described in non-genetically based PHAL tracing studies (36, 38), with the exception of a few weak projections (Table S1). Quantitative analysis of relative projection strength to each of 30 principal targets, using fluorescence pixel intensity, revealed strong projections within the hypothalamus, including the MPOA, PVN AHN, DMH and PMv, and the extended amygdala (including the BNST), as well as posterior projections to the midbrain and brainstem, including the PAGdm, PAGvl, MRN, and VTA (Fig. 1F, see color code for their position relative to VMHvl). A comparison of projections in males and females (n=2 each) did not reveal consistent differences (Table S2; but see (27).

**Figure 1.**
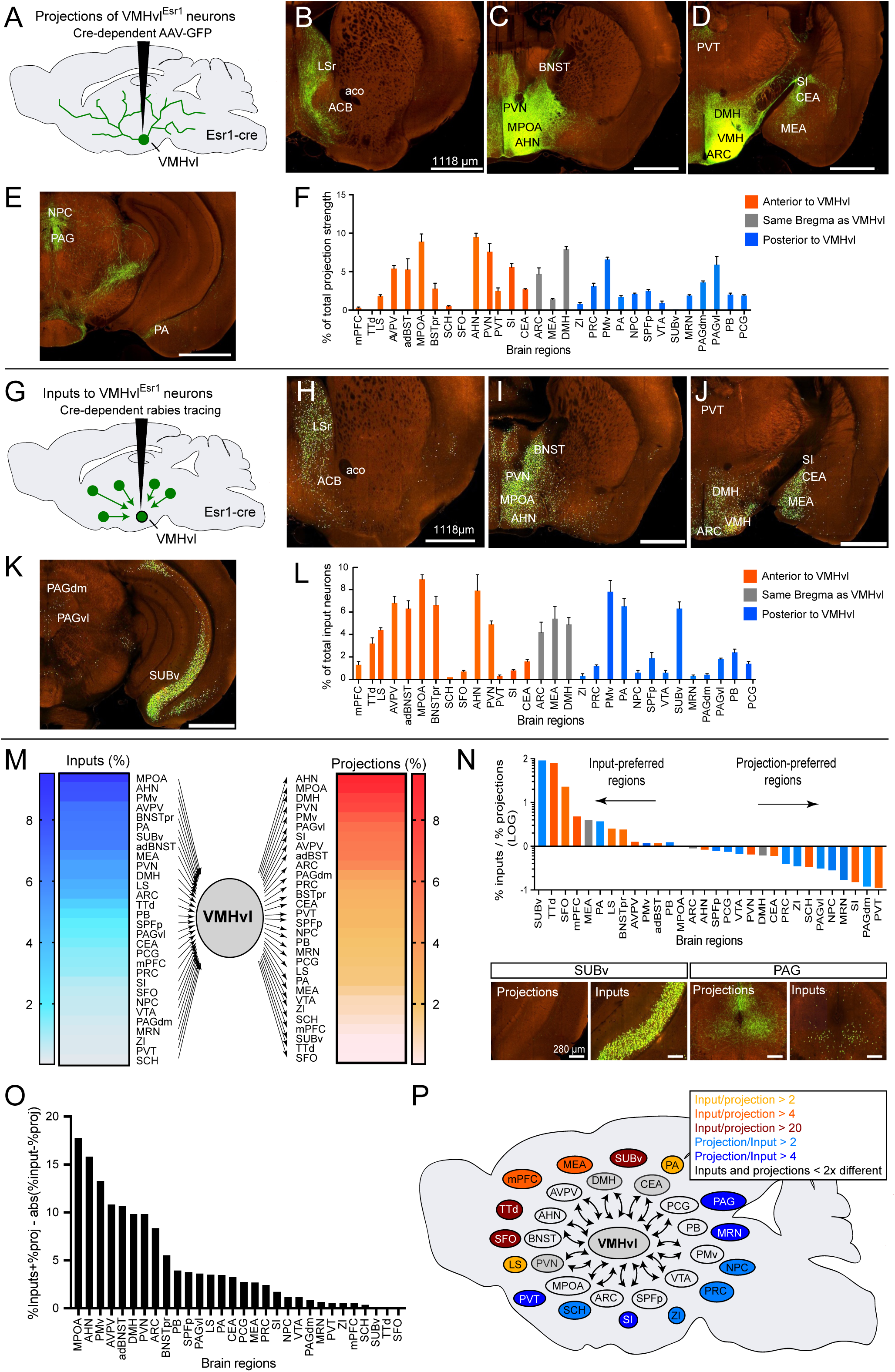
Inputs to and outputs from VMHvl^Esr1^ neurons. (A-F) Projection patterns of VMHvl^Esr1^ neurons. (A) Experimental schematic. (B-E) Representative images of AAV-GFP labeled projections (Image credit: Allen Institute). (B) from experiment #264319363, image #44. (C-E) from experiment #176886958, images# 63, #75, #88, respectively. (F) Quantification of proportion of total projection strength in each of 30 areas analyzed (n=3). Values in this and all other bar graphs with error bars represent means+SEMs. Bar graph color code corresponds to anterior (orange), posterior (blue), or similar (grey) A-P location (Bregma) relative to VMHvl; regions arranged from anterior (left) to posterior (right). See Table 1 for list of abbreviations and Table S4 for rank-ordering of input and projection strengths. (G-L) Inputs to VMHvl^Esr1^ neurons revealed by Cre-dependent, monosynaptic rabies tracing. (G) Experimental schematic. (H-K) Representative images of rabies-GFP labeled inputs. (L) Quantification of proportion of total inputs, graphed as in panel (F). (M) Schematic summarizing the percent of total inputs (left) or outputs (right). Heat map coloration corresponds to percent of total labeling. (N) Relative bias towards inputs or projection strength for each region, calculated by dividing numbers in L (normalized input strength) by F (normalized projection strength). Bottom panels show examples of structures with strong input (SUBv) or projection (PAG) bias. (O) Histogram for visualizing estimated strength and balance of reciprocal connectivity for each region analyzed, calculated by subtracting the absolute value of the difference between the data in L (normalized input strength) and F (normalized projection strength) from their sum (normalized input strength + normalized projection strength). (P) Schematic showing the relative bias of regions towards input and output using heat-map coloration and the high rates of recurrent connectivity, particularly within the hypothalamus. See Figure S1 for control experiments, micrographs of input and output regions, and additional data. See Table S5 and Table S6 for rank-ordered list and relative positions of regions.

**Table 1.** Structures analyzed in this study and their abbreviations. Reference table showing abbreviations used in the text and figures of this study (left) and their full names (right).

| <u>Abbreviation used</u> | <u>Full name</u> |
| --- | --- |
| mPFC | Medial prefrontal cortex |
| PL | Prelimbic area |
| ILA | Infralimbic area |
| TTd | Taenia tecta, dorsal part |
| LS | Lateral septal nucleus |
| adBNST | Bed nuclei of the stria terminalis, anterior-dorsal division |
| BNSTpr | Bed nuclei of the stria terminalis, principal nucleus |
| MPOA | Medial preoptic area |
| SCH | Suprachiasmatic nucleus |
| SFO | Subfornical organ |
| AHN | Anterior Hypothalamic nucleus |
| PVN | Paraventricular hypothalamic nucleus |
| SI | Substantia innominata |
| CEA | Central amygdalar nucleus |
| ARC | Arcuate hypothalamic nucleus |
| MEA | Medial amygdalar nucleus |
| DMH | Dorsomedial nucleus of the hypothalamus |
| ZI | Zona incerta |
| PRC | Precommissural nucleus |
| PMv | Ventral premammillary nucleus |
| PA | Posterior amygdalar nucleus |
| NPC | Nucleus of the posterior commissure |
| SPFp | Subparafascicular nucleus, parvicellular part |
| PIL | Posterior intralaminar thalamic nucleus |
| VTA | Ventral tegmental area |
| SUBv | Subiculum, ventral part |
| MRN | Mid brain reticular nucleus |
| PAGdm | Periaqueductal gray, dorsomedial area |
| PAGvl | Periaqueductal gray, ventrolateral area |
| PB | Parabrachial nucleus |
| PCG | Pontine central gray |

To map pre-synaptic inputs specifically to the VMHvl^Esr1^ population, we performed Cre-dependent monosynaptic, retrograde, trans-neuronal tracing (48, 49), using a modified rabies virus with enhanced cell viability (Yao et al., *in preparation*), which encodes a nuclear GFP reporter (Fig. 1G, Methods). Sections were analyzed using brain-wide serial 2-photon tomography, as above, and the number of cells in each region was manually quantified. We observed detectable retrograde labeling in 63 different structures, all but 7 of which were identified in an earlier non-genetically based retrograde tracing study from VMHvl in the rat using CTB (37) (Table S3). Notably, the vast majority of inputs to VMHvl^Esr1^ neurons are located subcortically, particularly in the hypothalamus and extended amygdala. However, we also detected significant input from the ventral hippocampus, as well as from the medial prefrontal cortex (Fig. 1H-L; Fig. S1B).

Strikingly, with a few notable exceptions (see below), mono-synaptic retrograde labeling was found in the majority of structures to which VMHvl^Esr1^ neurons also projected (Fig. 1F, L, MP; Fig. S1A, B). As many of these input regions themselves contain Esr1^+^ neurons, we were concerned that the Cre-dependent, trans-complementing AAV encoding TVA-2A-Rabies Glycoprotein (RG) injected into VMHvl might infect axons from these VMH-extrinsic Esr1^+^ inputs, subsequently allowing for primary infection of those terminals by the EnvA-pseudotyped rabies virus (and therefore GFP expression in their cell bodies). To control for this, a separate cohort of mice were injected with a control Cre-dependent AAV encoding only TVA, and not RG. With the exception of local expression of GFP in VMHvl, expression of the rabies-encoded nuclear GFP marker was not detected in input regions (Fig. S1C vs. D), indicating that the VMH-extrinsic GFP expression observed in experimental animals was dependent on the expression of the trans-complementing RG protein, and therefore indeed reflected trans-synaptic, retrograde tracing.

To compare quantitatively the relative strength of VMHvl^Esr1^ projections vs. inputs for each structure, we determined the ratio, plotted on a log scale, of the normalized input strength to the normalized output strength for 30 different regions (Fig. 1N, Table S4, S5). Thus, the closer the log ratio value to 0, the higher the likelihood that the normalized strength of inputs from a given region was equal to the normalized projection (output) strength to that region (e.g., MPOA, ~9% of total projections and ~9% of total inputs; ratio=1; log ratio=0; Figs. 1F, L-N). Approximately 40% (12/30) of the regions analyzed in this way had log ratio values between +0.3 and -0.3 (Fig. 1N), indicating relatively balanced (less than 2-fold different) reciprocal connectivity. In contrast, only 20% (6/30) of the regions sampled had an input bias (ratio >3, log ratio>0.48), while a similar percentage showed an output bias (ratio<0.33, log ratio <-0.48). Most notable among input-biased areas was the ventral subiculum (SUBv), which contributed ~7% of all inputs but which received almost no output, while among output-biased areas the PAG received ~5% of all outputs but <1% of all inputs (Fig. 1F, L-N, P).

To analyze these data further, we ranked input-biased structures and output-biased structures based on the ratio of the normalized inputs/outputs for each structure (Table S5 and Table S6). Interestingly, input-biased structures were distributed primarily anterior to, or in the same A-P region as, the VMHvl (Table S6, cols 2, 3), and included sensory regions (TTd, MeA, BNSTpr), while projection-biased structures were distributed more posteriorly and included most of the midbrain premotor targets (e.g., PAGdm, PAGvl, MRN) (Table S6, cols 5, 6). The average distance from Bregma was significantly different (p<.03) for input-biased vs. output-biased targets (Table S6, cols 4, 7, 10), with the former containing the most anterior (mPFC, TTd, LS) and the latter the most posterior (PAGdm, PAGvl, MRN) targets. To further analyze unbiased regions that have the greatest combination of both connectivity strength and recurrence, we plotted the sum of each region’s normalized inputs and projections (strength) minus the absolute value of the difference between input and projection (bias; Figure 1O). These strongly reciprocally connected regions included most intra-hypothalamic and extended amygdala targets (e.g., MPOA, AHN, BNST; Fig 1O, P; Table S6, cols 8, 9, 11, 12).

We next asked whether different input regions send predominantly glutamatergic or GABAergic projections to the VMHvl. To do this, we performed <u>i</u>nput-<u>n</u>eurotransmitter-<u>se</u>lective <u>r</u>etrograde <u>t</u>racing (INSERT) by injecting retrograde HSV harboring a Cre-dependent mCherry reporter into the VMHvl of either vGlut2-cre or vGAT-cre mice. Retrograde transport of the virus and recombination revealed the brain-wide distribution of vGlut2^+^ or vGAT^+^ neurons that project to the VMHvl (Fig. S1E), although not to Esr1^+^ neurons specifically. These experiments indicated that many (63%) of the VMHvl input regions are either primarily glutamatergic or GABAergic, but not both (Fig. S1F-H). The PA, PRC, SPFp, and PMv provide the majority of predominantly glutamatergic input, while the BNSTpr, MPOA, AHN, and DMH provide the majority of predominantly GABAergic input (Fig. S1F-H). Among those input regions containing both glutamatergic and GABAergic neurons, most send inputs of both types to the VMHvl. MPOA, AVPV, and ARC send equal proportions of glutamatergic and GABAergic inputs, while other regions show either a strong bias to GABAergic (BNSTpr and aTU) or to glutamatergic inputs (mPFC, MEAav, pPA, SUBv, and PAGdm) (Fig. S1F-H).

### Higher order projections from VMHvl reveals indirect feedback to select presynaptic inputs

To identify second and higher-order projections of VMHvl^Esr1^ neurons, we performed poly-synaptic, anterograde, trans-neuronal tracing using a Cre-dependent H129 virus (Fig. 2A) (50). Survival times were limited to ~40-50 hours to restrict viral movement to an estimated 2 ± 1 synapses (50, 51). As expected, post-synaptic labeling was seen in most of the structures to which VMHvl^Esr1^ neurons project directly, including the PAG, MPOA, ARC, and BNSTpr (Fig. 2B-I). Strikingly, however, strong labeling was also observed in several areas to which VMHvl^Esr1^ neurons do not directly project (Fig. 2I-L), but from which they receive strong input; these included the SUBv, TTd, and the SFO (Fig. 2L; S2E-G). These observations are suggestive of polysynaptic feedback from VMHvl to these input structures (Fig. 2L, Fig. S2J). (However we cannot formally exclude that this reflects input-specific retrograde labeling by the H129 virus, although this has not been described.) We also identified indirect outputs to the SCH (Fig. S2A-C), a structure that has recently been shown to mediate circadian control of aggression via indirect projections to VMHvl^Esr1^ neurons (52). Finally, although VMHvl^Esr1^ neurons exhibit very weak direct projections to mPFC, (Fig. 1F, N), we observed clear anterograde trans-synaptic labeling of cells in PL and ILA (Fig. 2B).

**Figure 2.**
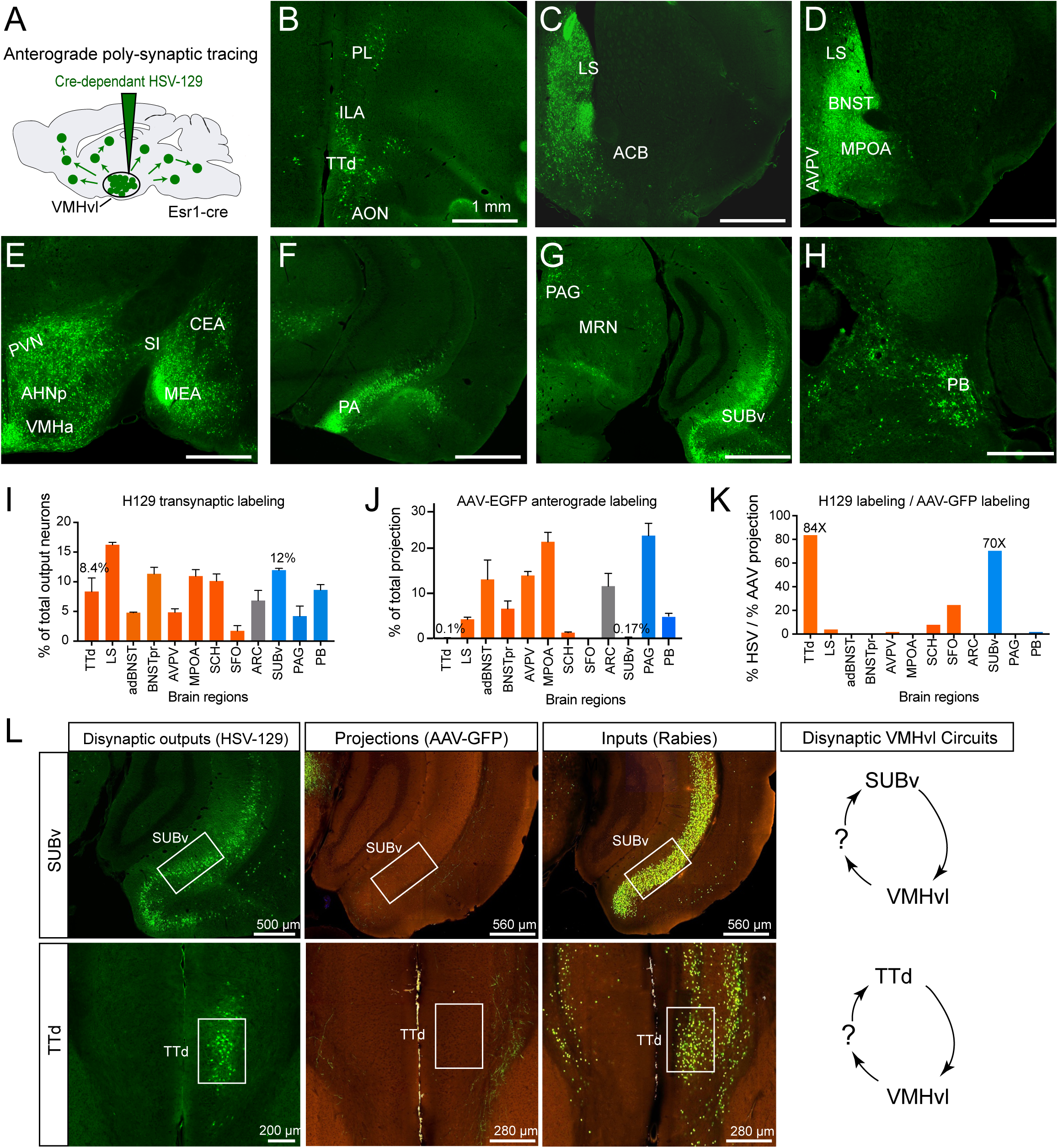
Indirect feedback loops revealed by anterograde trans-synaptic tracing. (A) Schematic of experiment showing anterograde, poly-trans-synaptic tracing via injection of H129 Δ TK-TT virus into the VMHvl of Esr1-cre mice. (B-H) Representative images of anterograde trans-synaptic tracing from VMHvl^Esr1^ neurons. (I) Quantification showing the percent of total H129 Δ TK-TT labeled output neurons in 12 regions (mean+SEM, n=2). Bar color code reflects whether the region is located anterior (orange) or posterior (blue) to VMHvl. (J) Quantification of AAV-GFP-based projections for the same 12 selected regions extracted from Figure 1F and normalized to 100% for comparison to data in panel I. (K) Ratio of H129 Δ TK-TT labeling to AAV-GFP projections (i.e. data from panel I divided by data from panel J). (L) Representative images comparing HSV labeled polysynaptic outputs, AAV-GFP-based projections, and rabies-GFP labeled inputs in the SUBv and TTd. Schematics of indirect feedback loops from VMHvl^Esr1^ to TTd and SUBv shown on the right. A di-synaptic connection is indicated for illustrative purposes but the data do not exclude poly-synaptic connectivity. See Figure S2 for additional data and analysis.

### VMHvl^Esr1^ neurons show extensive collateralization

The population-level projection mapping described above (Fig. 1) did not distinguish whether the multiple projection targets of VMHvl^Esr1^ neurons (Fig. 1F; S1A) reflect collateralization by individual neurons, or rather a mixed population, each of which has relatively restricted, specific projections. To investigate this issue, we performed collateralization mapping of VMHvl^Esr1^ neurons, using a modification of a previously described method (13). In this approach, a Cre-dependent, retrogradely transported HSV encoding FLPo recombinase is injected into one projection target of Esr1^+^ neurons, while a FLPo-dependent AAV encoding GFP is injected into the VMHvl of *Esr1-Cre* mice (Fig. 3A, B). This intersectional strategy results in GFP expression specifically in VMHvl^Esr1^ neurons that project to the target region of interest. Because GFP is freely diffusible in the cytoplasm, it fills each neuron, labeling any collateral projections to other targets, which can then be identified by GFP expression (Fig. 3A, B). This method is unbiased and more efficient at identifying collateral projections than is conventional dual-color retrograde labeling with CTb variants, which can only interrogate a few targets per animal. In this case, we used a Cre-dependent HSV-FLPo, rather than a Cre-dependent CAV-FLPo as originally reported (13) because we found that the latter exhibited a high degree of bias (tropism) for different types of inputs to the injected region.

**Figure 3.**
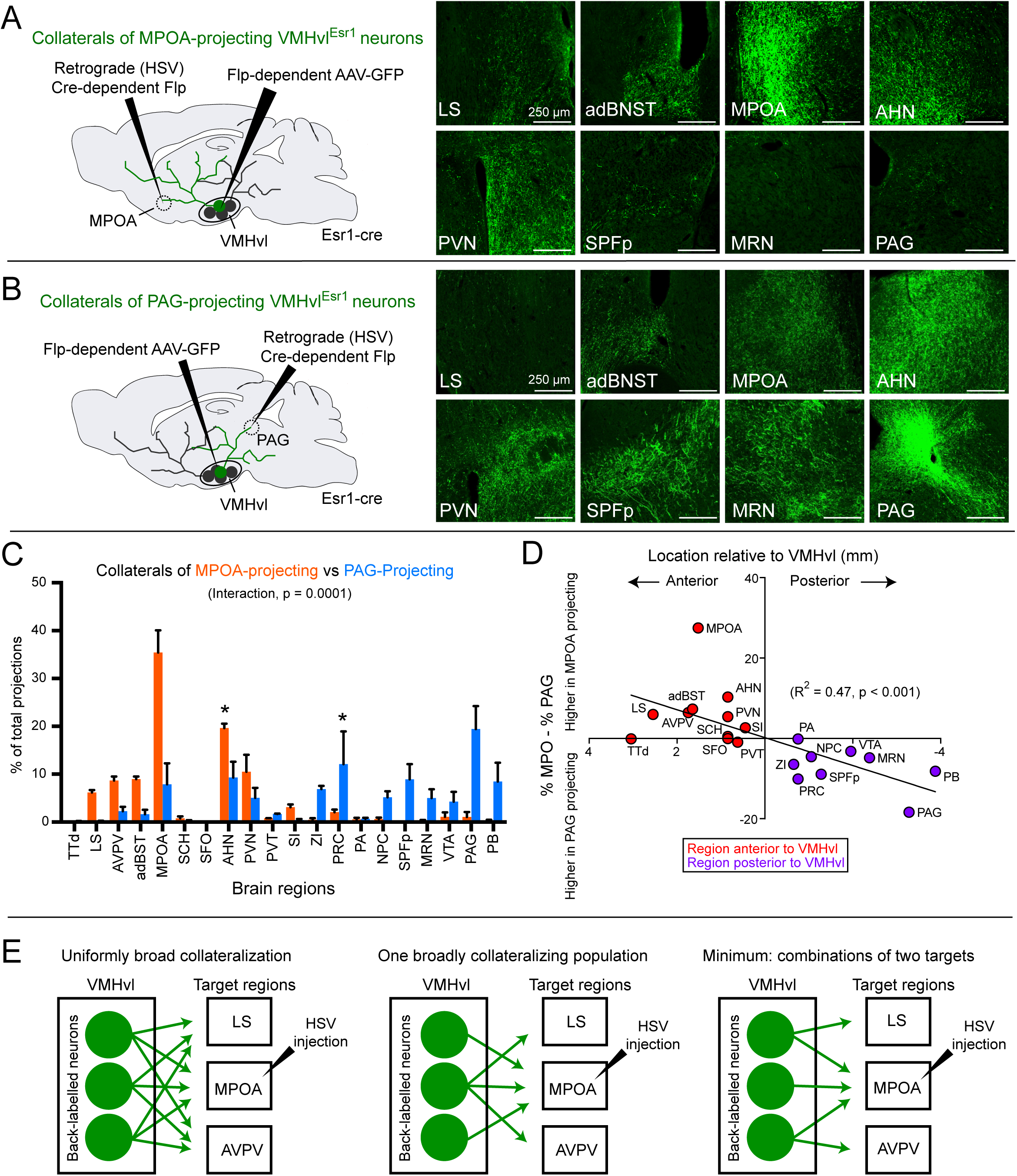
VMHvl^Esr1^ subpopulations collateralize to multiple, distinct targets. (A) Schematic illustrating experimental strategy for labeling collaterals of VMHvl^Esr1^ neurons that project to the MPOA. Representative images of coronal sections showing AAV-GFP expression in collateral projections of VMHvl^Esr1^ neurons that project to the MPOA shown on the right. (B) Schematic illustrating experimental strategy for labeling collaterals of VMHvl^Esr1^ neurons that project to the PAG. Representative images of coronal sections showing AAV-GFP expression in collateral projections of VMHvl^Esr1^ neurons that project to the PAG shown on the right. (C) Percent of total projection strength in each of 20 collateral targets for VMHvl^Esr1^ neurons that project to the MPOA (orange bars, n=2) and VMHvl^Esr1^ neurons that project to the PAG (blue bars, n=3). Bars show mean+SEM. There is a strong interaction effect by 2-way ANOVA (p=0.0001), with the AHN and PRC reaching significance after correcting post-hoc tests for multiple comparisons (p<0.05, Sidak correction). Projections to MPOA and PAG were omitted from the analysis. (D) Scatter plot of the location along the A-P axis relative to the VMHvl (x-axis) versus the difference in percent of total collaterals observed in each region between MPOA- and PAG-projecting populations (percent of total for MPOA – percent of total for PAG; y axis). There is a significant correlation between AP position and bias towards collateralization from MPOA (anterior regions) vs PAG (posterior regions) (R^2^ = 0.47, p < 0.001). (E) Schematics illustrating extreme possibilities for the underlying collateral connectivity that could yield these results. Individual neurons could target all downstream regions (left), there could be a single broadly collateralizing population (center), or, minimally, subpopulations could target different combinations of two downstream regions. See Figure S3 for collateralization analysis of LSv-projecting VMHvl neurons.

Initially, we injected Cre-dependent HSV-FLPo in two major targets of VMHvl^Esr1^ neurons, representing the anterior and posterior projection domains, respectively: MPOA and the dorsal PAG (Fig. 3A, B). In striking contrast to results reported for ARC Agrp^+^ and MPOA Gal^+^ neurons, we observed a high degree of collateralization among VMHvl^Esr1^ neurons, more reminiscent of midbrain neuromodulatory systems (13, 26, 53). For example, injection of HSV-LS1L-FLPo into the MPOA resulted in collateral projections to at least 5 major areas in addition to the MPOA itself, including the LS, AVPV, adBNST, AHN, and PVN (Fig. 3A, C). Similarly, injection of the retrograde virus into the dorsal PAG resulted in collateral projections to at least 6 different areas in addition to dPAG, including the adBNST, MPOA, AHN, ZI, SPFp, and MRN (Fig. 3B, C). From this we infer that at least some individual VMHvl^Esr1^ neurons collateralize to multiple (>1) targets (Fig. 3E).

### Subsets of VMHvl^Esr1^ neurons exhibit an anterior or posterior bias in collateralization pattern

Further analysis of these data revealed an unexpected finding: although there was qualitative overlap in the collateral targets of VMHvl^Esr1^ neurons back-labeled from either MPOA or dmPAG, there were important quantitative differences as well (Fig. 3C, D). VMHvl^Esr1^ neurons that project to MPOA, which is located anterior to VMH, collateralized significantly more strongly to other anteriorly located targets, while VMHvl^Esr1^ neurons projecting to dmPAG, which is located posterior to VMH, collateralized more strongly to other posteriorly located targets (Fig. 3C, D). 2-way ANOVA revealed a strong interaction between HSV^LS1L-FLPo^ injection site (MPOA versus PAG) and the distribution of labeling in collateral projection domains (Fig. 3C, p = 0.0001, MPOA and PAG omitted from the analysis). We also observed a strong correlation between the A-P location of target regions and their relative collateralization strength between MPOA- and PAG-projecting populations (Fig. 3D, R^2^ = 0.47, p<0.001).

A similar bias was observed when comparing the collateral projections of VMHvl^Esr1^ neurons back-labeled from LSv, another region anterior to VMH, to those back-labeled from the PAG (Fig. S3A-C), while comparison of collaterals of VMHvl^Esr1^ neurons back-labeled from LSv and MPOA revealed no significant differences (Fig. S3D-E). Thus, although VMHvl^Esr1^ neurons as a whole collateralize extensively, we find evidence for subpopulations that project to combinations of target regions located either anterior or posterior to the VMH itself (Fig. 3C, D). However, within these two subpopulations, these data do not distinguish between a range of possible connectivity patterns of the individual neurons (Fig. 3E).

### Projection-defined subpopulations of VMHvl^Esr1^ neurons differ in soma location and size

Our finding of projection-biased subpopulations led us to analyze characteristics of these Esr1^+^ subpopulations in VMHvl, particularly as previous studies have described differences in cell morphology in the VMH (54-56). Interestingly, we observed that the cell somata of the anteriorly vs. posteriorly projecting VMHvl^Esr1^ neurons described above exhibit a distinct and complementary distribution along the anterior-posterior axis: anteriorly-projecting cells (injected with retrograde HSV^LS1L-FLPo^ virus in MPOA or LSV) were located in the more posterior domain of VMHvl (-1.58 to -2.06 mm from Bregma), while posteriorly-projecting cells (injected with retrograde HSV^LS1L-FLPo^ virus in PAG) were located in the more anterior region of VMHvl (-1.34 to -1.82 mm from Bregma; Fig. 4A, B and Fig. S3F). This between-animal comparison was confirmed by a within-animal comparison using dual injections of HSV-LS1L-mCherry into MPOA and of HSV-LS1L-eYFP into the dmPAG of *Esr1-Cre* mice (Fig. 4C-D). Serial sectioning revealed opposing gradients of dmPAG- and MPOA-projecting neurons along the anterior-posterior axis, with the former peaking more anteriorly (-1.34 to -1.46 mm from Bregma) and the latter peaking more posteriorly (-1.7 mm from Bregma; Fig. 4D). Moreover, within individual sections containing both back-labeled populations, dmPAG-projecting and MPO-projecting VMHvl^Esr1^ neurons were largely non-overlapping (Fig. 4C, “overlay”, 4D; overlap less than expected based on random distribution, p<.01).

**Figure 4.**
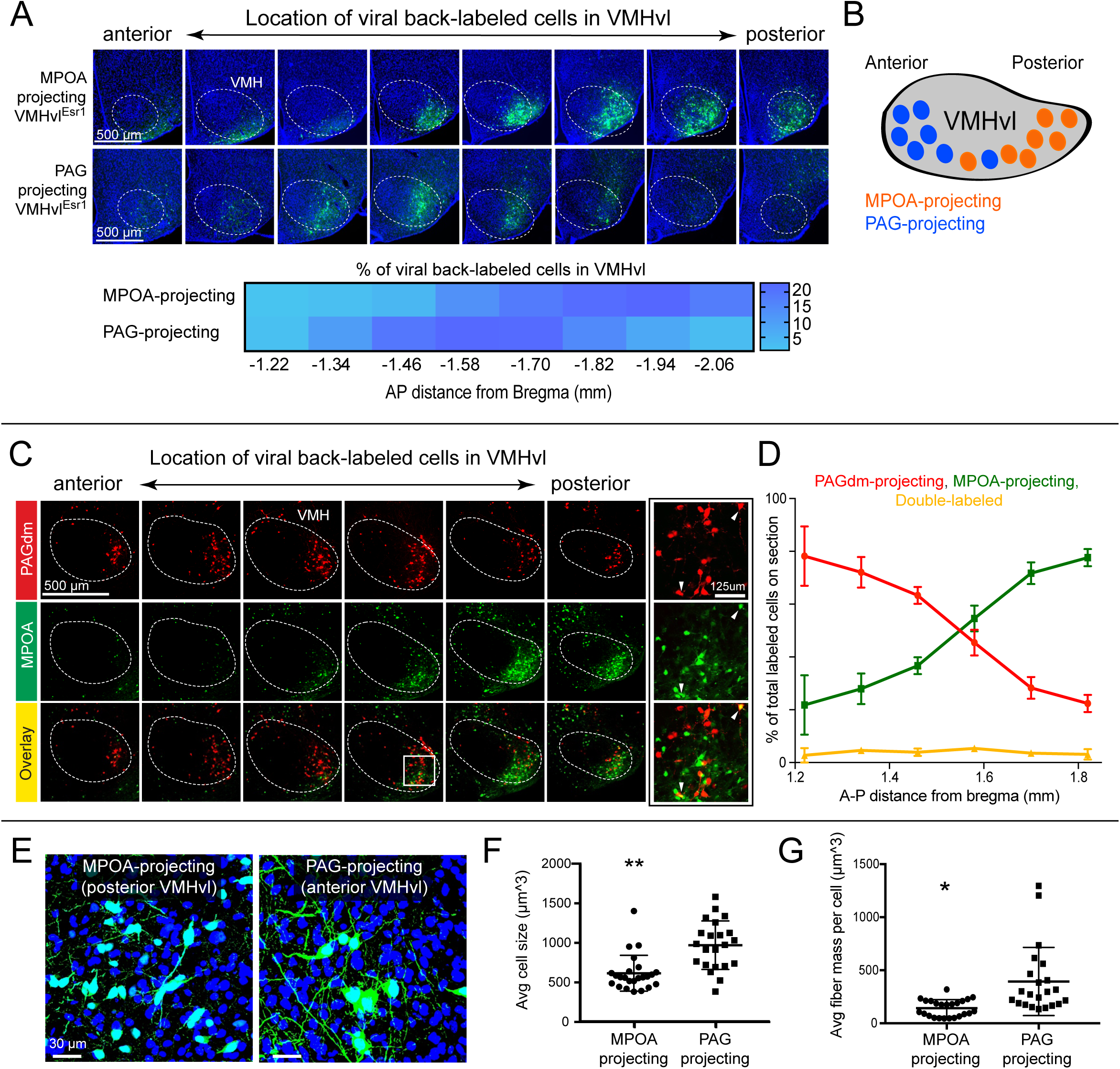
Distinct characteristics of VMHvl^Esr1^ neurons that project to the MPOA vs PAG. (A) Representative images (top) and heatmap (bottom) showing the distribution along the A-P axis of VMHvl^Esr1^ neurons back-labeled with Cre-dependent HSV that project to the MPOA (top) versus the PAG (bottom) (n=2/each). (B) Schematic illustrating the complementary position of back-labeled neurons in VMHvl. (C) Representative images of coronal sections showing the distribution of neurons in the VMHvl that were back-labeled via injection of Cre-dependent HSV-mCherry into the PAG and Cre-dependent HSV-EYFP into the MPOA of Esr1-cre mice. High magnification images of labeled VMHvl neurons shown in the rightmost panels correspond to the boxed region in the bottom-center micrograph. Arrowheads point to double-labeled cells. (D) Quantification of the distribution of back-labeled cells in the VMHvl (mean±SEM, n=3). (E) Representative confocal images of VMHvl ^Esr1^ cell bodies back-labeled using Cre-dependent HSV from either the MPOA (left) or PAG (right). (F) Cell volume of MPOA- vs. PAG-projecting VMHvl^Esr1^ neurons back-labeled as in E (mean±SEM, n=4 mice/each; total neurons: n=997 from MPOA and n=522 from PAG, p < 0.001). (G) Average fiber mass for MPOA- vs. PAG- projecting VMHvl^Esr1^ neurons back-labeled as in E (mean±SEM, n=4 mice/each, p <0.01).

Further analysis indicated that the average somata diameters of the MPOA- and PAG-projecting neurons were significantly different, with the latter being almost 60% larger than the former (964±65 μm^3^ vs. 609±46 μm^3^, p < 0.001; Fig. 4E, F). In addition, the average fiber mass per cell (an indirect measure of branching within VMHvl) was significantly larger for PAG-projecting than for MPOA-projecting Esr1^+^ neurons (p < 0.01; Fig. 4E, G). Taken together, these data reveal significant differences in cell body distribution (along the A-P axis of VMHvl), cell somata size, and local branching for anteriorly- vs. posteriorly-projecting Esr1^+^ neurons in the VMHvl, as well as minimal overlap among these cells at A-P positions where both are located, suggesting that they represent distinct subpopulations of Esr1^+^ neurons.

### Projection-biased subsets of Esr1^+^ neurons receive similar inputs

The existence of different subpopulations of Esr1^+^ neurons exhibiting anterior- or posterior biases in their collateralization patterns raised the question of whether these subpopulations differed in their inputs. To address this question directly, we performed a modification of the TRIO method (13), in which mono-synaptic, retrograde, trans-synaptic tracing is performed on subsets of Esr1^+^ neurons as defined by their projections. In this experiment, Esr1-Cre mice were injected in either the MPOA or PAG with a Cre-dependent HSV-LS1L-FLPo virus in order to express FLPo in Esr1^+^ neurons with anteriorly vs. posteriorly biased projections, respectively; these same mice were injected in the VMHvl with FLPo-dependent AAVs encoding TVA-mCherry and RG for rabies tracing. Following three weeks of incubation, the animals were additionally injected in VMHvl with EnvA-pseudotyped rabies^ΔRG^ expressing GFP (Fig. 5A, C). Following a further 6 days of incubation, animals were sacrificed and the pattern of GFP labeling quantified.

**Figure 5.**
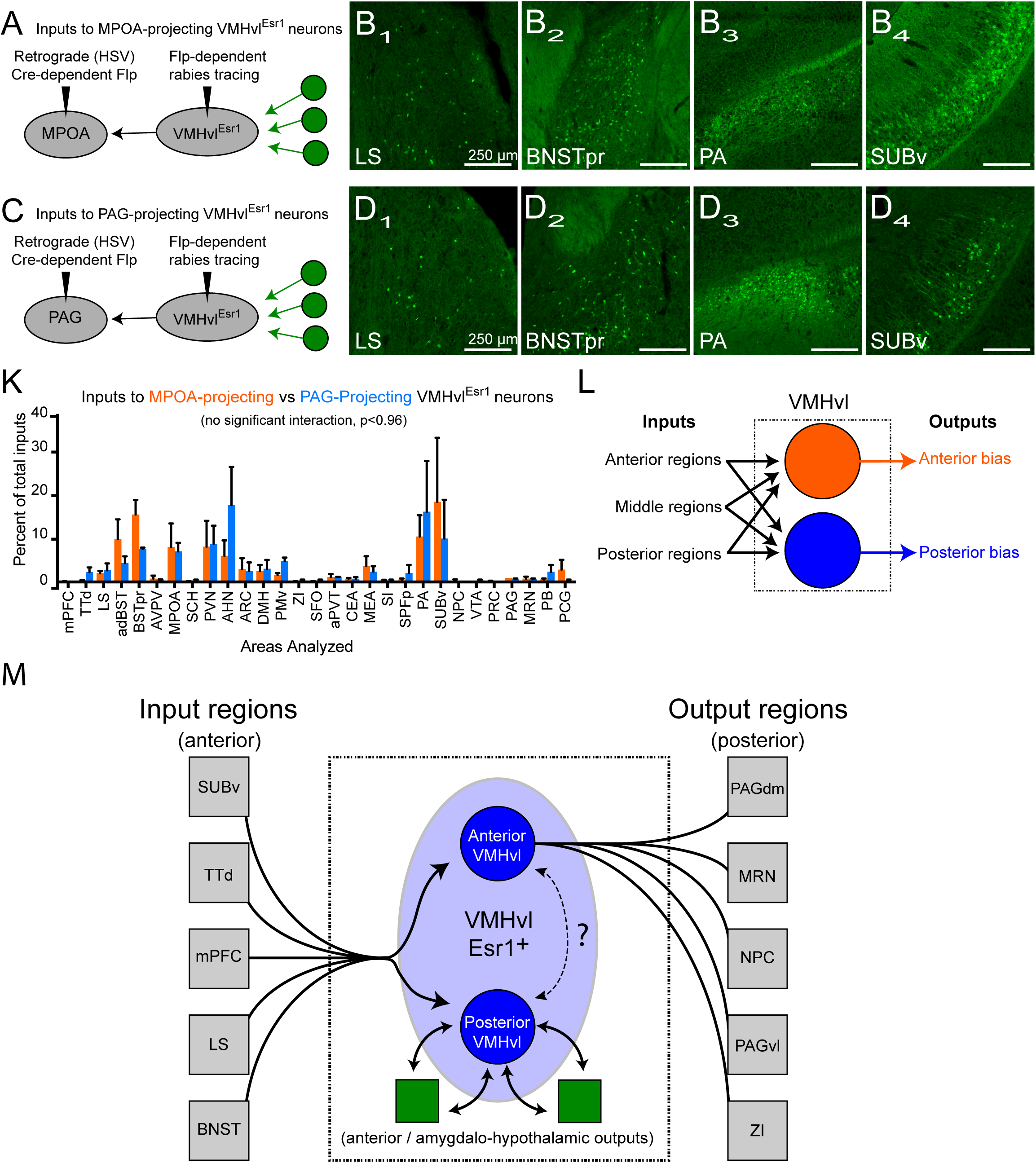
Projection-biased VMHvl ^Esr1^ subpopulations receive similar inputs. (A, C) Schematic of experiment for tracing inputs based on projection pattern. (B, D) Representative images of coronal sections showing rabies labeled inputs to VMHvl^Esr1^ neurons that project to the MPOA (B) versus to VMHvl^Esr1^ neurons that project to the PAG (D). (K) Quantification of the proportion of total inputs from 29 regions (mean+SEM, n=2/each). There is no significant interaction observed by 2-way ANOVA (p<0.96) or differences between specific regions. See Fig. S4 for distribution of soma locations for MPOA vs PAG back-labeled cells. (L) Schematic illustrating results of collateralization (Figure 3) and input-output mapping in which subpopulations of VMHvl^Esr1^ neurons collateralize to distinct combinations of anterior or posterior targets yet receive the same inputs. (M) Schematic showing overall summary of VMHvl^Esr1^ circuit architecture.

Interestingly, despite the differences in cell body location, size, and the separation of projection targets between these two VMHvl^Esr1^ populations, retrogradely labeled input cells could be found in similar structures following rabies tracing from each of these two VMH populations (Fig. 5B, D, K, L). There was no statistically significant differences in the proportion of back-labeled cells in each of 29 input areas analyzed, and no significant interaction between projection target and the proportion of labeling in input regions (2-way ANOVA, p<0.96, Fig. 5K). This does not exclude the existence of subtle but biologically significant differences that are obscured by the multiple sources of variance in this 4-virus experiment. To confirm that rabies labeling was in fact generated from the distinct projection populations of VMHvl^Esr1^ neurons described above (Fig. 4), we quantified the distribution of viral back-labeled and rabies-infected starter cells (Rabies-GFP^+^, TVA-mCherry^+^) along the anterior-posterior axis of the VMHvl in mice where rabies tracing was performed from PAG-back-labeled vs. MPOA-back-labeled cells. Similar opposing gradients of cell soma distributions were observed, as in the collateralization-mapping experiments (Fig. 4A), with the distribution of MPOA-back-labeled cells shifted posteriorly relative to that of PAG-back-labeled cells (Fig. S4). Together these data indicate that MPOA-projecting (posteriorly located, anteriorly collateralizing) vs. PAG-projecting (anteriorly located, posteriorly collateralizing) VMHvl^Esr1^ neurons receive a similar combination of presynaptic inputs (Fig. 5L).

## DISCUSSION

Esr1^+^ neurons in VMHvl control social investigation, mounting, and fighting as well as internal states related to these behaviors. To investigate the anatomical input-output logic of this brain region and to provide a foundation for understanding this functional diversity, we have carried out systematic mapping of inputs to and outputs from these neurons using 6 different types of viral-genetic tracing systems: 1) anterograde projection mapping; 2) retrograde, mono-synaptic, trans-neuronal tracing; 3) input-neurotransmitter-selective retrograde tracing (INSERT); 4) anterograde, polysynaptic, trans-neuronal tracing; 5) collateralization mapping; 6) mapping the relationship between inputs and projection patterns. To our knowledge, this is the first time that all of these techniques have been applied to a single, genetically identified cell population in the same study. This combination of tracing techniques has afforded a brain-wide overview of the collateralization patterns of Esr1^+^ neuronal projections, and of the input-output logic of these neurons as defined by the intersection of gene expression and projection pattern.

### Fan-in/fan-out organization and recurrence of VMHvl inputs and outputs

Previous studies of VMH connectivity using non-genetically targeted tracers have demonstrated that this nucleus receives inputs from and projects to diverse structures (37, 38). The present studies confirm and extend these and other classical studies of VMHvl connectivity (36, 41) in several important ways. First, they demonstrate a similarly high degree of input convergence and of output divergence for *specifically* the Esr1^+^ subset of VMHvl neurons. It is somewhat surprising that the overall pattern of inputs and outputs is so similar for Esr1-restricted vs. non-genetically restricted labeling of VMHvl neurons, given that the former only represent ~40% of the latter (7). However, the VMHvl^Esr1^ population is itself heterogeneous (9), and our data reveal heterogeneity in collateralization patterns within this population. Furthermore, some of the inputs and projections may specifically map to Esr1+ neurons in the subjacent tuberal (TU) region due to spillover of injected viruses.

Second, VMHvl^Esr1^ neurons demonstrate a high degree of recurrent connectivity between their inputs and outputs: of 30 input regions analyzed, 90% also receive feedback projections from Esr1^+^ neurons in VMHvl, and the remaining 10% receive indirect, polysynaptic feedback projections. Such a direct comparison would be difficult to make based on earlier studies, which typically examined either inputs or outputs, but not both. Notably, the vast majority of the inputs and outputs of VMHvl^Esr1^ neurons are located subcortically, particularly in the hypothalamus and extended amygdala. However, there is input from the ventral subiculum (hippocampus), as well as from the medial prefrontal cortex. Interestingly, the ventral hippocampus does not receive direct feedback from the VMHvl, although we observe an indirect recurrent projection to SUBv.

Third, using a two-virus collateralization mapping technique not previously applied to the VMH, we find that Esr1^+^ neurons collateralize to multiple (≥5) targets (“fan-out”). These findings contrast with previous studies of collateralization patterns in the nearby, hypothalamic ARC and MPOA: two predominantly GABAergic structures reported to project to different targets via independent, projection-specific cell populations (22, 24). Whether this reflects differences in methodology, function, or in the logic of GABAergic vs. glutamatergic projections in the hypothalamus, is an interesting question for future studies.

Fourth, and most important, we find that VMHvl contains at least two subpopulations of Esr1^+^ neurons with different cell body sizes, local branching patterns, and differential soma distributions along the A-P axis, which exhibit distinct anterior vs. posterior biases in their pattern of collateralization. Nevertheless, these two projection-defined, Esr1^+^ subpopulations of VMHvl neurons receive a similar distribution of “fan-in” inputs from the same anatomical regions, as assessed using the TRIO method (Fig. 5M). This input-output organization appears distinct from those described thus far using these techniques in other hypothalamic nuclei (discussed below).

### VMHvl^Esr1^ neurons collateralize broadly yet specifically

Previous studies have shown that the projections of ARC^Agrp^ neurons, which control hunger, exhibit little collateralization and a high degree of target specificity, suggesting a 1-to-1 organization (22). Similar observations have recently been reported for MPOA^Gal^ neurons involved in parenting behavior (24). Interestingly, both of those populations are GABAergic. Here we show that VMHvl^Esr1^ neurons, which are largely glutamatergic, exhibit substantial collateralization in their projections. While these population data demonstrate that at least some VMHvl^Esr1^ neurons collateralize to multiple (>1) targets, they do not prove that individual neurons collateralize to all 5-6 targets, nor do they exclude that some neurons may project exclusively to single targets (e.g., the regions where HSV-FLPo was injected; Fig. 3E.) Collateralization could, in principle, impose temporal synchrony on targets of VMHvl^Esr1^ neurons and coordinate their activity. Given this collateralization, assigning specific functions to individual projections (e.g., by optogenetic terminal stimulation, (22, 57) will be technically challenging, due to back-propagation and collateral redistribution of action potentials from optogenetically stimulated terminals.

Although VMHvl Esr1^+^ neurons project to multiple targets, we nevertheless observed that subpopulations of VMHvl^Esr1^ neurons exhibit biases in their collateralization to different combinations of target regions. One pattern is biased towards more posterior structures, while the other is biased towards anterior structures. The neurons exhibiting these collateralization biases, moreover, exhibit differences in anterior-posterior location within VMHvl, differences in cell body size and degree of local branching, and are largely non-overlapping in dual-labeling experiments. This suggests that they may represent distinct cell classes or cell types, possibly corresponding to those described using classical Golgi staining (58). Clarification of this issue could be achieved using single-cell RNA sequencing in combination with retrograde labeling, FISH, and quantitative analysis of cell morphology.

### Overall system architecture of VMHvl^Esr1^ neurons

Although the results presented here indicate that VMHvl^Esr1^ neurons exhibit a high degree of connectional complexity, some broad features of system architecture can be extracted (Fig. 5M). First, despite the high level of recurrent connectivity (Fig. 1), there are a few areas that are strongly biased inputs to, or outputs from, these neurons. The former include TTd, an olfactory processing area; the SFO, an interoceptive circumferential organ; the ventral hippocampus (SUBv), and the mPFC. These areas tend to be located anterior to VMH. The latter include the PAG and MRN, which are pre-motor regions located posterior to VMH. In contrast, VMHvl^Esr1^ neurons exhibit the strongest level of recurrent connectivity with other hypothalamic (MPOA, AHN, ARC, DMH) and extended amygdalar (MeA, BNSTpr) regions, many of which are located at similar A-P levels as VMH (Fig. 1P).

This pattern of connectivity suggests that there is a sparse anterior-to-posterior flow of information through the VMH, from sensory/cognitive to pre-motor regions, respectively, with a network of highly recurrent connectivity interposed between these input and output processing streams (Fig. 5M). The high degree of recurrent connectivity between VMHvl^Esr1^ neurons and the surrounding amygalo-hypothalamic structures is suggestive of an analog controller, which might regulate behavioral decision-making and/or internal state. Given the limited set of premotor target structures of VMHvl^Esr1^ neurons, such a system may function as a circuit dedicated to the finely tuned, dynamic control of a small set of social behaviors and their associated internal states, including aggression, social investigation, and mounting.

This overall system architecture provides an attractive *a posteriori* rationalization for the two subpopulations of VMHvl^Esr1^ neurons identified here. On the one hand, the posteriorly projecting population conveys output from the putative central controller to premotor structures that control behavior (Fig. 5M, right side). Consistent with this notion, preliminary experiments suggest that posteriorly projecting Esr1^+^ neurons express higher levels of c-fos following aggression than do anteriorly projecting neurons (DWK and DJA, unpublished). On the other hand, the anteriorly projecting population engages with the highly recurrent intra-hypothalamic and amygdalar circuitry (Fig. 5M, center). This circuitry may control internal state, competition with opponent behaviors, and/or go/no-go decisions gating social behavior. Such an organization raises the question of how these different VMHvl^Esr1^ populations communicate with, and are coordinated with, each other. Communication could involve local interneurons within VMHvl (Fig. 5M, dashed arrows), while coordination could be achieved through the parallelization of inputs to both subpopulations. While clearly speculative, this system architecture makes a number of predictions that can be explored in future studies.

### Open questions

The methods used in this study leave open several important questions. First, we do not yet know the degree of neuronal subtype diversity among VMHvl^Esr1^ neurons, and its relationship to overall VMHvl connectional architecture. It is already clear from recently published work that the VMHvl contains subpopulations of neurons with social behavior functions distinct from those involved in aggression (9, 46, 59, 60). Further, preliminary single-cell RNAseq analysis suggests that there are at least 4 transcriptomically distinct subsets of VMHvl^Esr1^ neurons, as well as ~12 other Esr1^-^ VMHvl cell populations (DWK and DJA, unpublished). How these subpopulations are integrated into the overall system architecture of the VMHvl is not yet clear, leaving open the possibility that simplifying features and principles will emerge when the connectivity and function of these more specific subpopulations is eventually mapped.

Second, we do not know the cellular identities of the various inputs and outputs identified in this study. We have identified both glutamatergic and GABAergic neurons that project to VMHvl, but which of these cells are mono-synaptic inputs to Esr1^+^ neurons, and how these specific inputs relate to downstream targets, remains unknown. Similarly, we do not know the precise combinations of downstream regions targeted by individual VMHvl^Esr1^ neurons, or the identities of the cells that are post-synaptic to these projections. The latter is important because a given recurrent projection could mediate either net positive- or net negative-feedback, depending on whether the immediate post-synaptic targets are, for example, excitatory or local inhibitory neurons, respectively. Connectivity mapping strategies involving trans-synaptic tracing, scRNAseq, and bar-coding (61) may help to resolve these issues.

Finally, and most importantly, a major frontier lies in understanding VMHvl function in the context of the broader meso-scale networks described here. Next-generation tools for simultaneously recording from and manipulating multiple brain regions simultaneously at cellular resolution, while integrating anatomic and transcriptomic information, offer exciting opportunities to clarify the fine-grained system architecture within this overall circuitry, and may reveal a simplifying organizational logic to the seemingly daunting complexity described here.

## ACKNOWLEDGEMENTS

We thank Dr. Miquel Chillón Rodriguez, Dr. Fan Wang, Dr. Liqun Luo and Dr. Rachael Neve for providing CAV-cre, retrograde Lenti-cre, TRIO reagents and retrograde HSV-fDIO-cre viruses, respectively, for pilot experiments; Dr. Lynn Enquist and the CNNV viral core center (Princeton University) for H129 recombinants, Dr. Kimberly Ritola of Janelia Research Campus for G-deleted EnvA rabies viruses, Bin Yang for help with cell-volume measurement software, Ben Ouellette for technical help, C. Chiu for laboratory management and G. Mancuso for administrative assistance. This work was supported by NIH grants 1R01MH070053 to D.J.A. and 1U01MH105982 to D.J.A. and Hongkui Zeng. B.W. is a Howard Hughes Medical Institute Fellow of the Life Sciences Research Foundation. D.J.A. is an Investigator of the Howard Hughes Medical Institute.

## AUTHOR CONTRIBUTIONS

LL designed and performed experiments, collected and analyzed data, made figures, and contributed to writing the manuscript, DK performed and analyzed data for duel labeling experiments, SY performed cloning experiments and produced the viral vectors for rabies experiments, AC designed and orchestrated production of rabies and associated viruses, JA orchestrated production of anterograde tracing data, HZ provided resources and support for AC and SY, DJA and BW supervised the project, contributed to data analysis, and wrote the manuscript.

## METHODS

### Animals and Viral Vectors

*Esr1^Cre/+^* mice (Lee et al, 2014) were generated and maintained at the California Institute of Technology (Caltech). vGAT-Cre and vGLUT2-Cre mice were obtained from Jackson Laboratory. All animal experiments were performed in accordance with NIH guidelines and approved by the Caltech Institutional Animal Care and Use Committee (IACUC). Viral vectors: (1) For the anterograde projection study, AAV1-CAG-FLEX-EGFP (titer ~8x10^12^ viral genomes(vg)/ml) were obtained from the University of Pennsylvania Gene Therapy Program. (2) For the retrograde input study, Cre-dependent AAV1-Syn-DIO-TVA-dTom-CVS-N2cG (titer ~4x10^13^ vg/ml), AAV1-Syn-DIO-TVA-dTom (titer ~4x10^13^ vg/ml), and pseudotyped, G-deleted rabies viruses (EnvA-CVS-N2c-histone-GFP, 5 x10^9^ infectious units (IU)/ml) were obtained from our collaborators at the Allen Institute for Brain Science (AIBS). (3) For the anterograde trans-neuronal tracing study, Cre-dependent HSV1-H129ΔTK-TT (Lo and Anderson 2011) was kindly provided by Lynn Enquist (Princeton University) with titer ~7 x10^9^ plaque-forming units (pfu)/ml. (4) For the collateral projection study, Cre-dependent retrograde HSV-hEF1-α-LS1L-mCherry-IRES-flpo HT (titer ~3 x10^9^ IU/ml) was obtained from Massachusetts Institute of Technology (MIT) McGovern Institute for Brain Research. AAVDJ-EF1α-fDIO-EYFP was obtained from the University of North Carolina at Chapel Hill Gene Therapy Center Vector Core (titer ~4 x10^12^ vg/ml). (5) For tracing inputs based on projection (TRIO, Schwarz et al., 2015), FLP-dependent AAVDJ-CAG-fDIO-TVA-mCherry (titer ~2 x 10^13^ vg/ml) and FLP-dependent AAV8-CAG-fDIO-RG (titer ~4 x10^12^ vg/ml) were obtained from the Stanford University Neuroscience Gene Vector and Virus Core. Pseudo-typed, G-deleted rabies EnvA-ΔG-B19-GFP (titer ~4x10^8^ transforming units (TU)/ml) were purchased from the Salk Institute Viral Vector Core. Pseudo-typed, G-deleted rabies EnvA-N2c-GFP (titer ~2 x 10^8^ IU/ml) were kindly provided by HHMI Janelia Virus Service Facility. Cre-dependent retrograde HSV-hEF1-α-LS1L-BFP-IRES-flpo HT for TRIO and Cre-dependent retrograde HSV- hEF1 -α-LS1L-mCherry/GFP for duel retrograde labeling were obtained from MIT McGovern Institute for Brain Research.

### Stereotaxic Surgery, Virus Injections, and immunohistochemistry

*Esr1^cre^* mice (8-12 weeks old, male and female) were anaesthetized with 3-5% isoflurane for induction and 1-3% for maintenance. Mice were mounted on a stereotaxic frame (David Kopf Instruments) with heating pad placed underneath. For anterograde tracing, AAV1-FLEX-EGFP viruses were injected by iontophoresis (3 μA at 7 s ‘on’ and 7 s ‘off’ cycles for 5 min total) with a pulled glass capillary (World Precision Company) into VMHvl^Esr1^. Region of interest (ROI) was positioned using Model 1900 Stereotaxic Alignment System (David Kopf Instruments). For VMHvl, coordinates are anteroposterior (AP): –1.5 mm from Bregma; mediolateral (ML): +0.78 mm from the midline, dorsoventral (DV): –5.775 mm. Mice were euthanized four weeks later with Ketamine (100 mg/kg) and Xylazine (10 mg/kg) and brains were analyzed by TissueCyte 1000 serial two-photon (STP) tomography system at AIBS.

For monosynaptic retrograde labeling, AAV1-DIO-TVA-dTom-N2cG (or AAV1-DIO-TVA-dTom for rabies glycoprotein-less control) viruses were injected by iontophoresis (3 μA at 7 s ‘on’ and 7 2 ‘off’ cycles for 5 min total) into the VMHvl of Esr1-cre male mice, followed by ~500 nl of rabies EnvA-N2c-histone-GFP with a Nanoliter 2010 injector (World Precision Company) three weeks later into the same location. Mice were euthanized 9 days later and brains were analyzed by TissueCyte 1000 serial two-Photon (STP) tomography system at AIBS.

For the glutamatergic and GABAergic inputs study, ~200 nl of HSV-LS1L-mCherry virus was injected into the VMHvl of vGLUT2-cre or vGAT-cre male mice (same coordinates as above, except ML= –0.78 for these and for injections described below). Mice were euthanized ~3 weeks later via perfusion with 4% PFA and serial sections were generated using a cryostat. For poly-synaptic anterograde tracing, ~150-300 nl of HSV1-H129ΔTK-TT viruses were injected into the VMHvl of Esr1-cre male mice with a Nanoliter injector and mice were euthanized between 40 to 48 hours after injection depending on assessments of their health. Ketoprofen (5 mg/kg, SQ) was given each day until euthanization. To visualize tdTomato encoded by HSV1-H129ΔTK-TT tracers, sections were stained with rat anti-mCherry (Thermo Fisher Scientific, clone 16D7, Cat# M11238, RRID:AB_253661, 1:200) overnight at 4 C, followed by Alexa-488-donkey anti-rat (Invitrogen, 1:200) at room temperature for 3 hours. For the collateral projection study, ~200 nl of HSV-LS1L-flpo viruses were injected into projection targets of VMHvl^Esr1^ neurons of Esr1-cre male mice (LSv coordinates, AP= +0.14, ML= –0.55, DV= -4; PAG coordinates, AP= -3.88, ML= -0.137, DV= –2.375; MPOA coordinates, AP= +0.02, ML= -0.3, DV= -5.375). At the same time, ~200 nl of FLP-dependent AAVDJ-fDIO-EYFP was injected into the VMHvl. Mice were perfused with 4% PFA 28-30 days after injection and serial sections were made using a cryostat.

For the projection-based input study (modified TRIO, Schwarz et al., 2015), ~100-150 nl of HSV-LS1L-flpo viruses were injected into the projection targets (MPOA or PAG) of VMHvl^Esr1^ neurons in Esr1-cre male mice. At the same time, a 1:1 mixture of AAVDJ-fDIO-TVA and AAV8-fDIO-RG viruses were injected into the VMHvl (~150 nl total). Three weeks later, rabies EnvA-ΔG-GFP was injected into the same location. Mice were euthanized 6 days later via perfusion with 4% PFA and serial sections were generated using a cryostat.

For duel-retrograde labeling, 200 nl of HSV-LS1L-mCherry and 200 nl of HSV-LS1L-EYFP was injected into the PAG and MPOA, respectively, in the same VMHvl^Esr1^ male mouse. Mice were euthanized ~3 weeks later via perfusion with 4% PFA and serial sections were generated using a cryostat.

In all experiments, we targeted the Esr1+ cluster of the VMHvl, which forms a continuous population with the neighboring Tuberal nucleus (TU). The average % of viral backlabeled cells in these two regions combined was 77% from 8 injected mice selected from across experiments. The remaining labeling could be found in other nearby regions that express Esr1, particularly the ventral premammillary nucleus (PMv) and Arcuate Hypothalamic nucleus (ARC).

### Image Acquisition and Analysis

For projection and input studies, through collaborations with AIBS, brain samples with GFP-labeled axonal projections or rabies-labeled cells were imaged using the TissueCyte 1000 serial two Photon (STP) tomography system (0.35 μm *x-y* resolution and 100 μm z-sampling interval) (Ragan et al, 2012, Oh et al., 2014). For quantification of projection strength or input neuron number, ROIs were selected and screenshots taken at the same magnification. Projection mapping images are publicly available and can be found at (http://connectivity.brain-map.org). Experiments used in the projection study are Experiments #176886958, #264248605, 264319363 posted on AIBS website (© 2015 Allen Institute for Brain Science. Allen Brain Atlas API. Available from: brain-map.org/api/index.html). Both Allen Mouse Brain Atlas (2004) and Allen Mouse Brain Connectivity Atlas (2011) were used in this study. Thirty ROIs were chosen based on (1) projection or input strength, (2) relative position, located anteriorly or posteriorly, to VMHvl, (3) research interests in psychiatric disorders and innate behaviors.

For anterograde trans-neuronal tracing, collateral projections, glutamatergic vs. GABAergic inputs (INSERT), and projection-based TRIO, images were acquired by confocal microscopy for starter cell images (Oympus FluoView FV1000), or by slide scanner for the quantification of starter cell location and labeling outside of the VMHvl (Olympus VS-ASW-S6). For quantification, matching images of ROIs were obtained using the “Crop to Clipboard” function of the slide scanner software. Twenty-eight ROIs (glutamatergic or GABAergic inputs), twenty ROIs (collateral projection), twelve ROIs (H129 transsynaptic anterograde tracing), and twenty-nine ROIs (TRIO) were chosen based on the criteria listed above. Additionally, for H129 labeling, SUBv, TTd, SFO, and SCH were chosen as we observed strong polysynaptic labeling. For quantification of projections, images were analyzed using Fiji/ImageJ by manually setting a threshold (common to all images) to the point that tissue autofluorescence was not visible followed by identification of objects using particle analysis (“analyze particles” function), binarization of the image, and summation of the total positive area (i.e. the area that is covered by projections above threshold). For quantification of input/output cell number, images were processed as described for projection analysis, except watershed-based identification of objects was used prior to particle analysis which was used for object counting, with size and circularity limited to an appropriate range to identify individual cells. Automatic particle counting was visually inspected for accuracy and manually adjusted. For all ImageJ analysis, normalized projection strength or inputs were taken from three adjacent sections 30 or 60 μm apart. For cell size and fiber mass quantifications, confocal images (60x) obtained using an Olympus FluoView 1000 confocal were analyzed using Imaris Image Analysis Software (Bitplane, Oxford Instruments). To obtain optical image quality, stacked 60x confocal images of VMVHvl neurons were saved as OIB files and 3-D reconstructed using Imaris processing software followed by automated identification and quantification of fluorescent objects based on manually identified criteria (cells: volume threshold 100 μm^^^3; fibers: volume threshold 10 to 100 μm^^^3).

